# Spread of pathological tau proteins through communicating neurons in human Alzheimer’s disease

**DOI:** 10.1101/555821

**Authors:** Jacob W. Vogel, Yasser Iturria-Medina, Olof T. Strandberg, Ruben Smith, Alan C. Evans, Oskar Hansson, for the Alzheimer’s Disease Neuroimaging Initiative, and the Swedish BioFinder Study

## Abstract

Tau is one of the two pathological hallmarks of Alzheimer’s disease, and bears a much closer relationship to local neurodegeneration and cognitive impairment than the other hallmark, *β*-amyloid. Cell and rodent models have shown evidence that tau spreads from cell to cell through anatomical neuronal connections, and that this process is facilitated by the presence of *β*-amyloid. We test this hypothesis in humans by using an epidemic spreading model (ESM) to simulate the spread of tau over human neuronal connections, and we compare the simulated pattern of progression to the observed pattern measured in the brains of 312 individuals on the Alzheimer’s disease spectrum, using PET. Fitting our model, we found that the majority of variance in the overall pattern of tau progression could be explained by diffusion of an agent through the human connectome, measured using either functional connectivity or diffusion tractography. These models far exceeded chance, and outperformed models testing the extracellular spread of tau over Euclidian space. Surprisingly, the ESM predicted the spatial patterns of tau irrespective of whether subjects demonstrated evidence for brain *β*-amyloid. In addition, in *β*-amyloid-positive subjects only, regions with greater amyloid burden showed greater tau than predicted by connectivity patterns, suggesting a role of amyloid in accelerating the spread of tau in certain isocortical regions. Altogether, our results provide strong evidence that tau spreads through neuronal communication pathways even in normal aging, and that this process is accelerated by the presence of brain *β*-amyloid.

## 1. Introduction

Alzheimer’s disease is characterized by the presence of *β*-amyloid plaques and neurofibrillary tangles of hyper-phospohrylated tau at autopsy. Both of these pathological phenomena can now be quantified spatially in the brains of living humans using positron emission tomography (PET), allowing for the study of disease progression before death and, indeed, before symptoms manifest [1]. *β*-amyloid plaques are detectable in the brain many years or even decades before dementia onset [2], but appear to have only subtle effects on cognition and brain health in humans [3, 4, 5, 6], if any. In contrast, tau neurofibrillary tangles are strongly correlated with local neurodegeneration and, in turn, cognitive impairment [7, 8]. However, tau tangle aggregation in the medial temporal lobes is a common and fairly innocuous feature of normal aging [9, 10, 11]. Frank cognitive impairment often coincides with the spreading of tau tangles out of the medial temporal lobes and into the surrounding isocortex, a process that animal models have suggested may be potentiated or accelerated by the presence of *β*-amyloid plaques [12, 13].

Due to its close link with neurodegeneration and cognitive impairment, tau has received special attention as a potential therapeutic target for Alzheimer’s disease [14]. Perhaps the most compelling features of tau pathophysiology are its rather focal distribution of aggregation and its highly stereotyped pattern of progression through the brain. Specifically, neurofibrillary tangles first appear in the transentorhinal cortex, before spreading to the anterior hippocampus, followed by adjacent limbic and temporal cortex, association isocortex, and finally to primary sensory cortex [15, 10, 16, 17]. This very particular pattern has led many to speculate that pathological tau itself, or a pathological process that incurs tau hyper-phosphorylation and toxicity, may spread directly from cell to cell through anatomical connections [18, 19]. Strong evidence in support of this hypothesis has come from animal models, which have repeatedly demonstrated that human tau injected into the brains of *β*-amyloid expressing transgenic rodents leads to the aggregation of tau in brain regions anatomically connected to the injection site [20, 21, 22, 23, 12]. An important caveat to the aforementioned studies is that they involve injection of tau aggregates that greatly exceed the amount of tau produced naturally in the human brain. In addition, the studies were performed in animals that do not get Alzheimer’s disease naturally.

Unfortunately, there are many obstacles to studying the tau-spreading hypothesis in humans. While autopsy studies have provided evidence for tau spreading [24, 25], this evidence comes in the form of limited snapshots in deceased individuals. Tau-PET allows for the quantification of tau *in vivo*, but the PET signal is contaminated by off-target binding that limit interpretations [26, 27, 28, 29]. Despite this limitation, circumstantial evidence has emerged supporting the hypothesis that tau spreads through connected neurons in humans. Studies decomposing the spatial distribution of tau-PET signal in the human brain have revealed spatial patterns highly reminiscent of brain functional networks [30, 31]. In addition, brain regions with greater functional connections to the rest of the brain tend to have greater tau accumulation [32], and correlations have been found between functional connectivity patterns and tau covariance patterns [33, 34].

Despite mounting evidence linking brain connectivity and tau expression, the aforementioned studies mostly involve either comparisons between coarse whole-brain measures of tau and brain connectivity, or are limited to only a fraction of brain connections. The initial seeding of tau in the cortex is thought to lead subsequently to secondary seeding events that cascade systematically through the cerebral cortex. Therefore, it is paramount that studies assessing the spread of tau through the brain can effectively model the complex spatio-temporal dynamics of this process. Therefore, we test the tau-spreading hypothesis by placing a “tau seed” in the entorhinal cortex, simulating its diffusion through measured functional and anatomical connections, and comparing the simulated pattern of global tau spread with actual pattern derived from tau-PET scans of 312 individuals. This method allows for a cascade of secondary tau seeding events to occur along a network over time, more closely simulating proposed models of tau spread in the brain. We then examine how the behavior of our model interacts with brain *β*-amyloid.

## 2. Materials and Methods

### 2.1. Participants

Participants of this study represented a selection of individuals from two large multi-center studies: the Swedish BioFinder Study (BioF; http://biofinder.se/) and the Alzheimer’s Disease Neuroimaging Initiative (ADNI; adni.loni.usc.edu). Both studies were designed to accelerate the discovery of biomarkers indicating progression of Alzheimer’s disease pathology. Participants were selected based on the following inclusion criteria: participants must i) have an AV1451-PET scan, ii) have either a *β*-amyloid-PET scan (for ADNI: [^18^F]-Florbetapir, for BioF: [^18^F]-Flutemetamol) or lumbar puncture measuring CSF *β*-amyloid1-42. In addition, participants were required to be cognitively unimpaired, have a clinical diagnosis of mild cognitive impairment, or have a clinical diagnosis of Alzheimer’s dementia with biomarker evidence of *β*-amyloid positivity. For both cohorts separately, PET-based *β*-amyloid positivity was defined using a previously described mixture modeling procedure [5]. For BioFINDER, *β*-amyloid1-42 positivity was defined as an (INNOTEST) level below 650ng/L [35]. All participants fitting the inclusion criteria with AV1451 scans acquired (BioFINDER) or that were available for public download (ADNI) in May 2018 were included in this study. In total across both studies, 162 cognitively unimparied individuals, 89 individuals with mild cognitive impairment and 61 amyloid-positive individuals with suspected Alzheimers dementia were included. Demographic information can be found in Table 1.

**Table 1:** Demographic information.

|  | **CN** | **MCI** | **AD** | **Total** |
| --- | --- | --- | --- | --- |
| **n** | 162 | 89 | 61 | 312 |
| **Age (SD)** | 72.0 (6.4) | 70.84 (7.8) | 72.0 (7.9) | 71.7 (7.1) |
| **% Women** | 45.1% | 64.0% | 58.6% | 53.1% |
| **Education (SD)** | 14.8 (3.6) | 15.3 (3.7) | 12.8 (3.9) | 14.6 (3.8) |
| **% ApoE4** | 41.9% | 58.4% | 68.5% | 51.7% |
| **% Amyloid Positive** | 42.6% | 64.0% | 100.0% | 66.2% |

CN = cognitively normal; MCI = mild cognitive impairment; AD =
Alzheimer’s disease dementia, SD = Standard Deviation

### 2.2. PET Acquisition and Pre-processing

MRI and PET acquisition procedures for ADNI (http://adni.loni.usc.edu/methods/) and BioF [36] have both been previously described at length. All AV1451-PET scans across studies were processed using the same pipleine, which has also been previously described [36, 31]. Briefly, 5-min frames were reconstructed from 80-100 minutes post-injection. These frames were re-aligned using AFNIs 3dvolreg (https://afni.nimh.nih.gov/) and averaged, and the mean image was coregistered to each subject’s native space T1 image. The coregistered image was intensity normalized using an inferior cerebellar gray reference region, creating standard uptake value ratios (SUVR).

### 2.3. Transformation of PET data to regional tau-positive probabilities

Mean regional tau-PET SUVRs were extracted from each individual’s native space PET image using the Desikan-Killiany atlas [37], an 83-region atlas based on structural morphometry. All cerebellar regions were removed from the atlas, leaving 78 regions in total. Previous AV1451-PET studies have noted considerable off-target binding of the AV1451 signal, leading to signal in regions without pathological tau burden, and likely to pollution of signal in regions accumulating tau [26, 27, 29, 31]. While many previous studies have ignored these issues, accounting for off-target binding is essential to the current study, as our model cannot distinguish off-target from target signal, and we are not interested in the propagation of off-target signal. To address this issue, we utilized regional Gaussian mixture modeling under the assumption that the target and off-target signal across the population are distinct and separable Gaussian distributions (Fig 1A).

**Figure 1:**
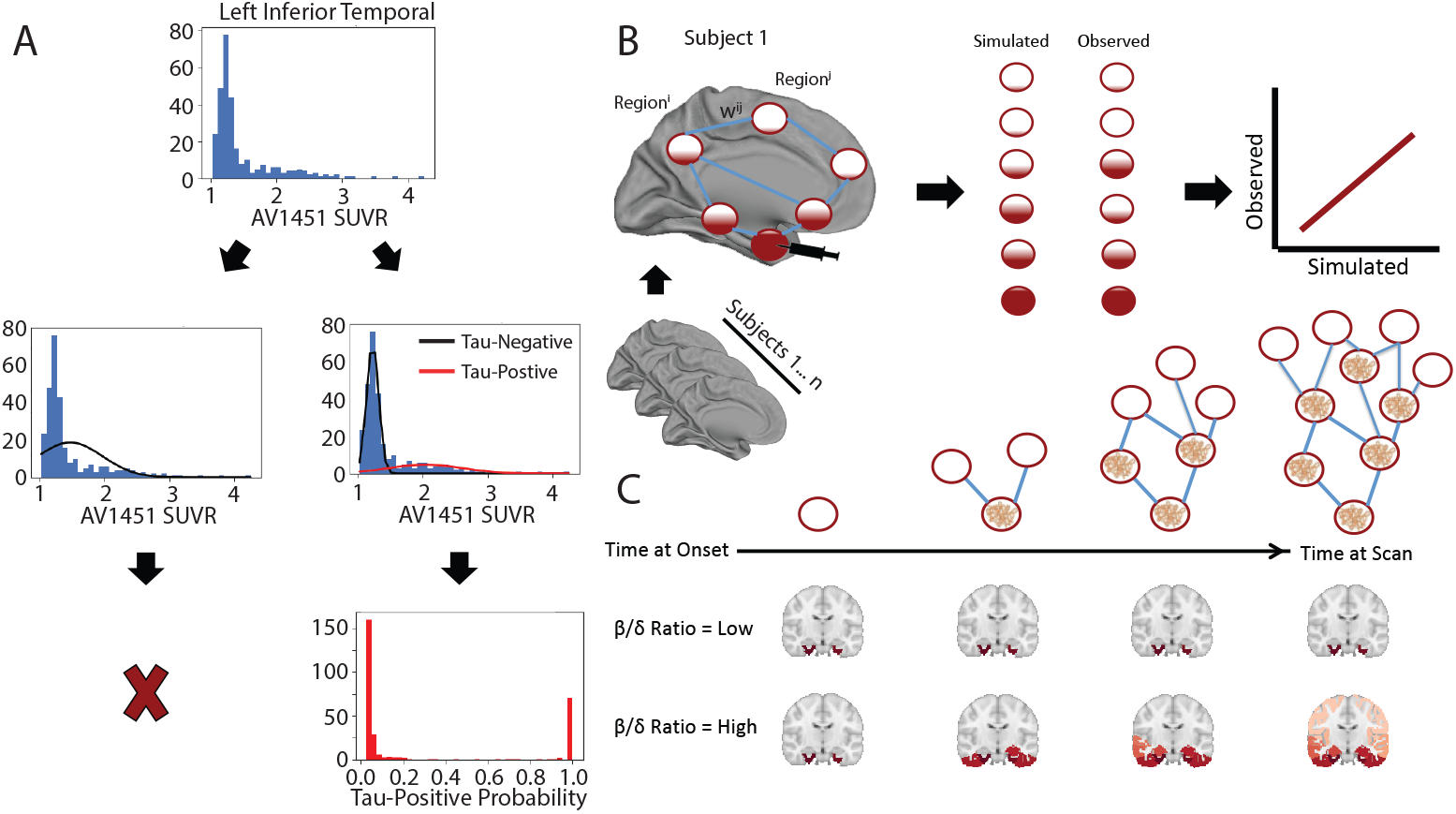
Methodological approaches. A) The distribution of all SUVR values in the left inferior temporal ROI are shown. Two Gaussian mixture models are fit to the data. When a one-component model fits the data better, the ROI is discarded. When a two-component model fits better, the probability that each values falls upon the second distribution is calculated. B) An artificial system based on a pairwise relationship (e.g. functional connectivity) matrix is created, where the relationship between regions i and j is represented by weight ij. For each subject, a seed is placed at the model epicenter, and the diffusion of this signal over time is simulated through the system, where the inter-regional relationships determine the pattern of spread, and subject-level free parameters determine the velocity of diffusion, until an optimal fit is reached. The simulated tau signal is then compared to the observed tau-PET signal to evaluate the model. C) Advantages of the ESM over traditional approaches includes the initiation of secondary seeding events as the diffusion process reaches new regions (top), and the fitting of subject-level production (*β*) and clearance (*δ*) parameters. A balance in these parameters will lead to little to no spreading over time, while increasing imbalance leads to accelerated spread.

As most individuals do not have tau in most regions, pathological signal should show a skewed distribution across the population, whereas off-target and non-specific signal should be reasonably normally distributed. Such a bimodal distribution has been observed for *β*-amyloid, and mixture modeling has been used in this context to define global *β*-amyloid positivity [38, 39]. Our approach differs from these previous studies as we do not assume the distribution of target and off-target binding to be homogeneous across cortical areas – we apply Gaussian mixture modeling separately to each region-of-interest (Fig 1A). Specifically, for each region, we fit a one-component and a two-component Gaussian mixture model across the entire population. We compare the fit of the two models using Aikake’s information criterion. If a two-component model fits the data better, this likely indicates the presence of pathological tau in a proportion of the population, and the Gaussians fit to the data provide a rough estimate of an SUVR threshold, above which AV1451 signal has a high probability of being abnormal. If a one-component model fits better, this indicates the AV1451-PET signal within the region is roughly normally distributed across the population, which we do not expect for tau in a population including many cognitively impaired individuals. Regions showing a unimodal distribution are therefore discarded from the ESM model, as neurofibrillary tau tangles are likely not expressed in that region within the sample. Furthermore, since the ESM receives regional (tau) probabilities as input, we calculate the probability that a given subject’s ROI SUVR value falls onto the second (i.e. right-most) Gaussian distribution using repeated five-fold cross-validation. Assuming this second distribution represents the subjects with abnormal AV1451 signal, this value estimates the proximity of a subject to the pathological distribution. Effectively, this converts regional SUVRs to regional tau-positive probabilities. This approach defines a fairly conservative, data-driven threshold for SUVR values, above which, one can assume the presence of abnormal signal (perhaps indicating pathological tau accumulation) with a high degree of confidence.

### 2.4. Connectivity measurements

The overall pattern of spread simulated by the Epidemic Spreading Model (see next section) is determined by the relationship matrix, which represents pairwise relationships between each region-of-interest. Indeed, this is the system through which the simulated signal will diffuse. Varying the relationship matrix can, for example, allow for the tests of different hypotheses of spread. We use a functional connectivity matrix generated from a group of young healthy controls to test the hypothesis that tau spreads through communicating neurons. We validate this procedure using anatomical connectivity measurements generated from healthy and impaired older adults. Finally, we test the hypothesis of tau spreading through extra-cellular space by using a Euclidian distance matrix as input.

Functional connectivity measurements were generated from a subsample of young healthy controls from the COBRE dataset [40], a publicly available sample which we accessed through the Nilearn python library. All subjects listed as healthy controls under the age of 40 were selected, totaling 74 individuals. The images were already preprocessed using the NIAK resting-state pipeline (http://niak.simexp-lab.org/pipepreprocessing.html), and additional details can be found elsewhere [40]. Correlation matrices were generated by finding the correlation between timeseries’ of each pair of regions-of-interest from the Desikan-Killiany atlas, and all available confounds were regressed from the correlation matrices. We took the mean of all 74 correlation matrices to create an average healthy connectome template. This connectome was then thresholded so as to only retain the top 10% of connections, and transformed so all values fell between 0 and 1.

To validate our findings, we created a template structural connectivity matrix using DTI tractography data from a non-overlapping sample of healthy and cognitively impaired individuals from ADNI. In total, 204 individuals had one or more DTI scans available, for a total of 540 scans. All scans were preprocessed with a previously described diffusion tractography pipeline [41], and acquisition and processing information has been described in detail [42]. Briefly, orientation distribution functions (ODF) were calculated and in turn used to generate deterministic connections between pairs of brain regions from the Desikan atlas. Specifically, an ACD measure was used, representing the total proportion of regional surface area (across both regions) that contain connecting fibers between the two regions. All images were assessed for quality. Connectomes were averaged across all subjects resulting in a template structural connectome in aging.

To create a Euclidian distance matrix, we calculate the coordinate representing the center of mass for each region of interest, and found the Euclidian distance between it and the center of mass of every other ROI. By using this distance matrix in the epidemic spreading model, we test the hypothesis that tau diffuses radially across adjacent cortex, rather than through connected regions.

### 2.5. The Epidemic Spreading Model

The spread of tau through connected brain regions was simulated using the Epidemic Spreading Model (ESM), a previously described diffusion model that has been applied to explain the spread of *β*-amyloid through the brain [43]. The ESM simulates the diffusion of a signal from an epicenter through a set of connected regions over time (Fig 1B,C). The dynamics of the spreading pattern are controlled by the weighted connectivity between regions, and by a set of parameters fit within-subject, the latter of which are solved through simulation. Specifically, the parameters represent subject-specific i) global tau production rate, ii) global tau clearance rate and iii) age of onset, which interact with regional-connectivity patterns to determine the velocity of spread. The ESM is simulated over time for each subject across several parameter sets, and the set that produces the closest approximation to observed tau burden for a given subject is selected. Note that these parameters themselves do not control regional patterning, which is the metric by which the accuracy of the model is evaluated (see below). Instead, the free parameters moderate the overall tau burden (i.e. the stopping point), which allows the ESM to be fit to individuals across the Alzheimer’s disease spectrum. For example, an individual with little-to-no tau burden would likely be fit with a balance of production and clearance rates that would preclude the overproduction and spread of tau signal (Fig 1C). A detailed and formalized description of the ESM can be found elsewhere [43].

In previous applications of the ESM, the model is fit over every possible epicenter as well as combinations of epicenters, and the epicenter providing the best overall fit to the data is selected. In our case, autopsy work provides strong evidence for a consistent “epicenter” of tau neurofibrillary tangles in humans. Tangles first emerge in the trans-entorhinal cortex, before emerging in other parts of the entorhinal cortex as well as the anterior hippocampus [15, 10]. We therefore ran models with the left and right entorhinal cortex selected as the model epicenters. However, for the purposes of validation, a best-fitting model-derived epicenter was also computed, by fitting the ESM across all possible regions and finding the best average within-subject fit. Once this epicenter was found, we ran the model once more using both left and right regions as the model epicenters.

The ESM takes as input a Region x Subject matrix of values ranging from 0 to 1, representing the probability of a pathological burden (in this case, of tau) in a given region for a given subject. The model is fit within-subject and, for each subject, produces an estimate of tau probability for every region-of-interest.

### 2.6. Experimental Design and Statistical Analysis

The ESM was fit using different relationship matrices (see above). Each model was evaluated by mean within-individual fit, as well as global population fit. Individual model fit is calculated as the r^2^ between predicted regional tau probabilities and actual regional tau probabilities measured with AV1451-PET, for each individual. The mean r^2^ across all individuals was used to represent overall model fit. To evaluate the accuracy of the global pattern, the regional predicted and observed tau probabilities, respectively, were averaged across all subjects, and the r^2^ between these group-averaged patterns were calculated. Together, these two accuracy measures represent the degree to which regional connectivity predicts the spatial pattern of tau-PET measured within and across subjects, respectively. To ensure the magnitude of our results were greater than chance given a matrix of similar properties, we fit the ESM using 100 null matrices with preserved degree and strength distributions using the Brain Connectivity toolbox (https://sites.google.com/site/bctnet/). We use the null distribution to calculate the mean and 95% confidence intervals of the relationship occurring by chance. Since we run only 100 null models per test, the lowest possible p-value is 0.01, which would suggest the observed test value was higher than all values observed by chance.

To examine the global accuracy of the ESM stratified by amyloid status, we first divided all subjects into one of two diagnostic groups: amyloid-negative and amyloid-positive. We then calculated the mean of predicted and observed values across all subjects within each amyloid group, respectively.

Studies in rodents have suggested a role of amyloid in facilitating the rapid fibrillarization of tau oligomers [12]. This would suggest that amyloid may play a role in explaining tau patterns that is at least partially independent of connectivity patterns. To explore this, we tested the relationship between regional modeling error and regional amyloid depositon. Amyloid-PET scans were available for 307/312 individuals, and were processed identically to tau-PET scans. We converted regional amyloid SUVR values to amyloid-positive probabilities using the same regional mixture-modeling approach as described above. Next, we used the sign of the residual to divide regions into those that were overestimated by the ESM, and those that were underestimated by the ESM. An underestimated region, for example, would show more tau than the model predicted given that region’s connectivity to the model epicenter. We explored the relationship between model estimation and amyloid by comparing the degree of (group-mean) amyloid between overestimated and underestimated regions using t-tests. We also observed this relationship within amyloid-negative and amyloid-positive subjects separately. In this case, the same (whole sample mean) amyloid measurements were used for both comparisons, but the regional under/overestimation varied by amyloid group.

## 3. Results

AV1451-PET scans measuring tau neurofibrillary tangles *in vivo* were available for 312 individuals spanning the Alzheimer’s disease spectrum. Demographic information for this sample can be found in Table 1.

### 3.1. Conversion to tau-positive probabilities enhances fidelity of tau-PET data

We executed a procedure to mitigate off-target binding of AV1451-PET data using mixture modeling. Regional Gaussian mixture modeling of AV1451 SUVR data across all 312 subjects suggested a two-component (bimodal) model as a superior fit for all 62 cortical regions-of-interest, as well as the left and right hippocampi and amygdalae. For all other subcortical regions-of-interest, a one-component model fit the data better, and these regions were discarded from all further analyses. The remaining 66 regions were converted to tau-positive probabilities (Fig 1A) using the Gaussian mixture models. This threshold-free, data-driven transformation yielded a sparse data matrix with a clear pattern suggesting a gradual progression of tau across regions of the brain (Fig 2). When sorted from least to most tau (e.g. [16]), the regional ordering greatly resembled the previously described progression of tau pathology [15].

**Figure 2:**
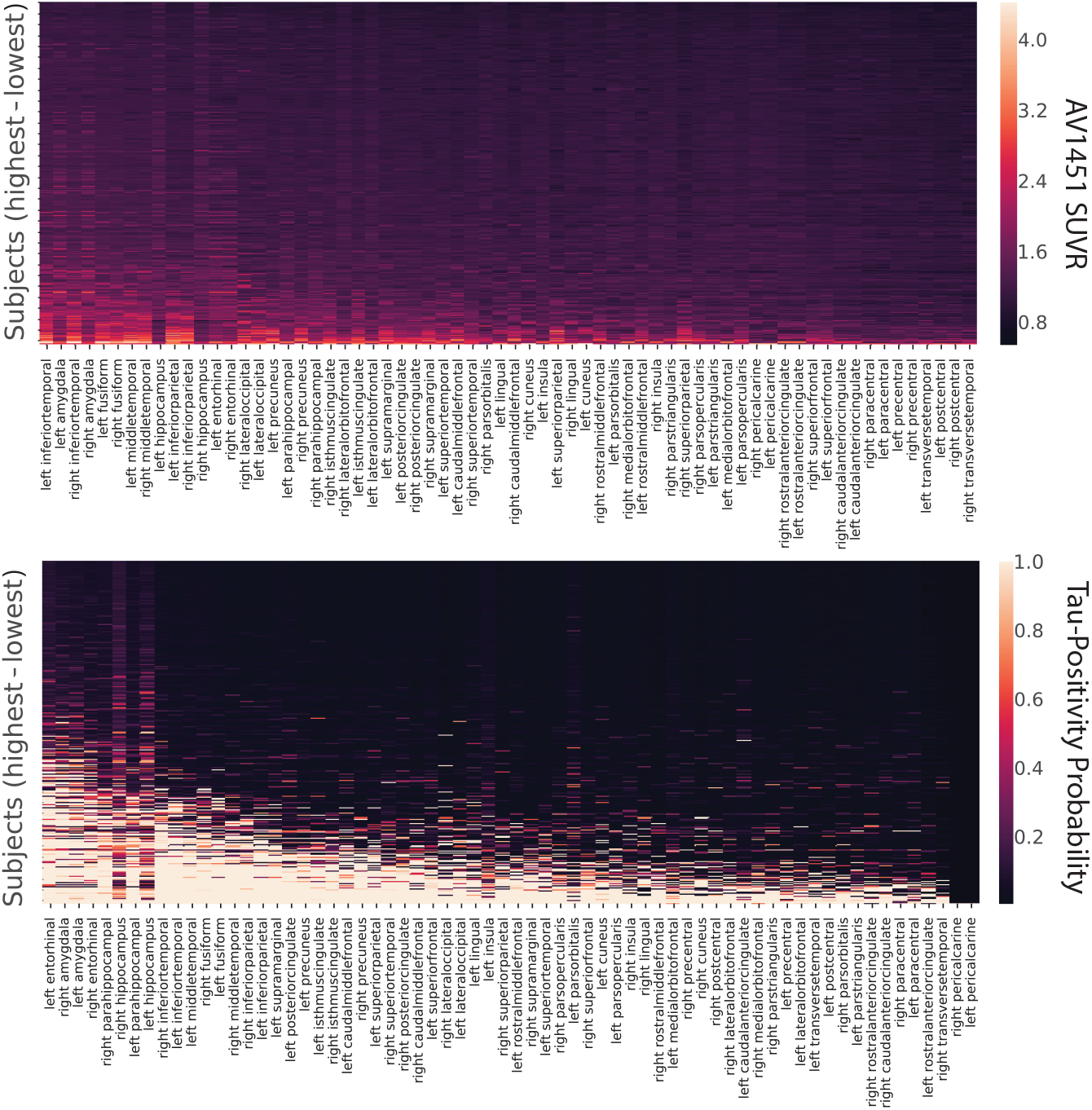
Tau-PET data before and after conversion to tau-positive probabilities. Each row is a subject sorted top-bottom by least to most overall tau. Each column is an ROI, sorted left to right by most to least overall tau. Warmer colors represent higher SUVR values (top) or tau-positive probabilities (bottom). Conversion to tau-positive probabilities creates a sparse distribution of values demonstrating a progression. The order of ROIs resembles those described in the autopsy literature.

### 3.2. Epidemic spreading of tau over human neuronal connections explains spatial pattern of tau in the brain

An epidemic spreading model was fit to the data, simulating the spread of tau signal from a single epicenter through functional brain connections over time (Fig 3,4). When using the left and right entorhinal cortex as the model epicenter, the model explained 59.6% (null model mean r^2^ [95% CI] = 0.060 [0.006, 0.126], p<0.01) of the overall spatial pattern of tau (Fig 4A), and on average, explained 33.6% (SD=20.0%; null model mean r^2^ [95% CI] = 0.068 [0.033, 0.147], p<0.01) of the spatial pattern within individual subjects.

**Figure 3:**
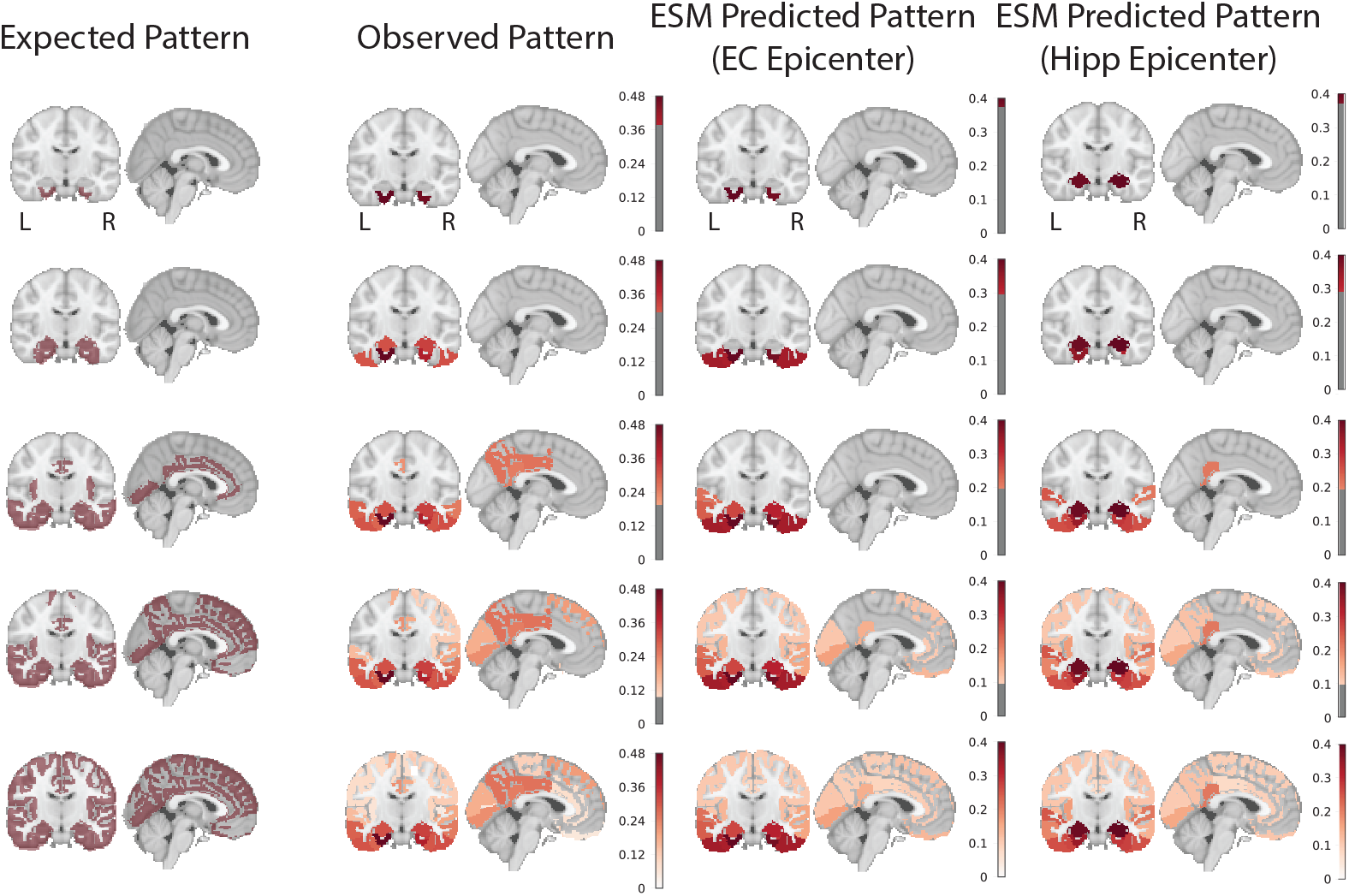
Hypothesized, observed and predicted pattern of tau spreading. (left) Hypothetical spread patterns represented by Braak stages I, II, VI, V and VI as described in [44]. (right) Spreading patterns of (from left to right) the observed tau-PET data, the ESM simulated data with entorhinal epicenter, and with hippocampus epicenter. Warmer colors represent higher proportion of regional tau-positivity predicted or observed across the population. Each “stage” was achieved by arbitrarily thresholding the population-mean tau-positive probability image at the following thresholds: 0.4, 0.3, 0.2, 0.1, 0

**Figure 4:**
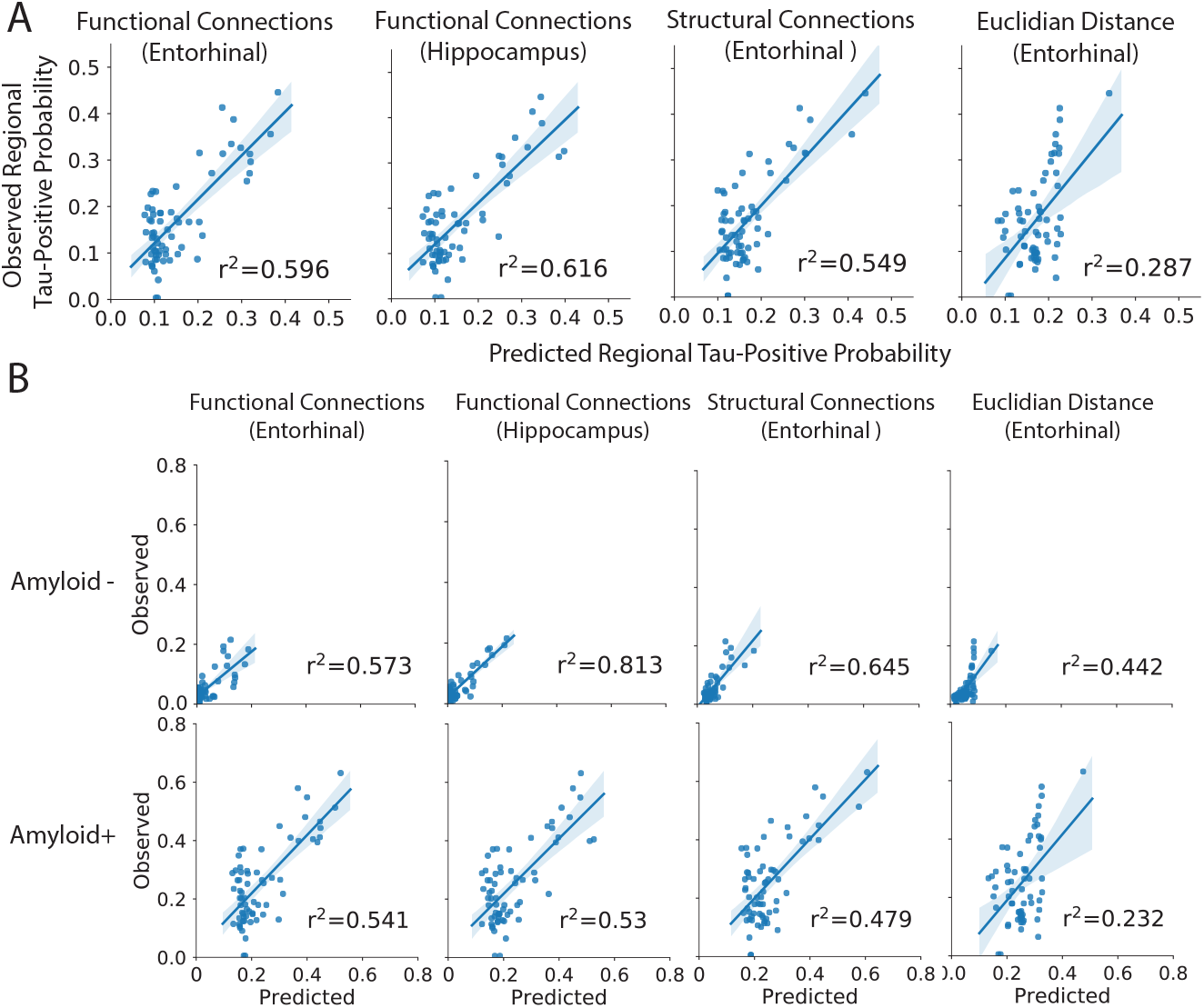
Performance of ESM in predicting spatial progression of tau. A) For each plot, each dot represents a region. The x-axis represents the mean simulated tau-positive probabilities across the population, while the y-axis represents the mean observed tau-positive probability. A value of (say) 0.3 for a given ROI would suggest that an average of 30% of all subjects included were predicted (X) or observed (Y) to have positive abnormal tau signal in that region. The results are shown for ESM fit over (from left to right) healthy functional connectome with entorhinal epicenter; healthy functional connectome with a hippocampus epicenter (selected as best-fitting); aging structural connectome with an entorhinal epicenter (also selected as best-fitting); and a Eucidian distance matrix with entorhinal epicenter. B) Breakdown of ESM performance by amyloid status. The average performance of the four different models are shown separately for amyloid-postive and amyloid-negative individuals.

Next, the ESM was fit allowing the model to select the “best-fitting” regional epicenter (Fig 4A). The hippocampus was selected, slightly improving the overall global accuracy of the model to 61.6%, but dramatically increasing the average local (within-subject) explained variance to 47.4% (SD=27.6%). The epidemic spreading model was particularly effective in predicting the early progression of tau, but diverged more from the observed tau pattern over time (Fig 3,4).

As a validation, the ESM was fit using a structural connectome created using diffusion tensor imaging tractography from a separate sample of healthy and cognitively impaired older adults (Fig 4A). The model fit was highly consistent with models fit over functional connectomes of younger adults. Using a bilateral entorhinal cortex epicenter, the model explained 54.9% (null model mean r^2^ [95% CI] = 0.062 [0.020, 0.133], p<0.01) of the overall spatial pattern of tau progression, and on average, explained 38.0% (SD=22.1%, null model mean r^2^ [95% CI] = 0.132 [0.108, 0.186], p<0.01) of the within-subject variance in tau spatial pattern. Once again, we fit the ESM allowing for a data-driven epicenter to be selected, and this time, the entorhinal cortex was selected as the best-fitting epicenter.

Alternative hypotheses have been proposed suggesting tau may simply spread extracellularly across neighboring regions, rather than through anatomical connections. To test this hypothesis, a model was fit over a Euclidean distance matrix instead of a functional or structural connectome (Fig 4A). This model explained considerably less variance, both at the global (r^2^=0.29) and individual (mean r^2^=0.23) level.

### 3.3. Low-level tau spreading is evident and predictable in amyloid-negative individuals

We divided our study sample into groups based on amyloid status and examined model accuracy separately within these groups. Model accuracy remained high even among amyloid-negative individuals despite a low overall tau burden (Fig 4B). This was validated by examining model fit against the tau pattern of individual amyloid-negative subjects (Fig 5). Model performance was high across most CN-subjects (Fig 5A), including those with low (Fig 5C) or even very low (Fig 5B) regional tau burden. In many cases, tau levels that would otherwise be considered sub-threshold nonetheless demonstrated a systematic pattern predicted by brain connectivity, particularly when using a hippocampal epicenter.

**Figure 5:**
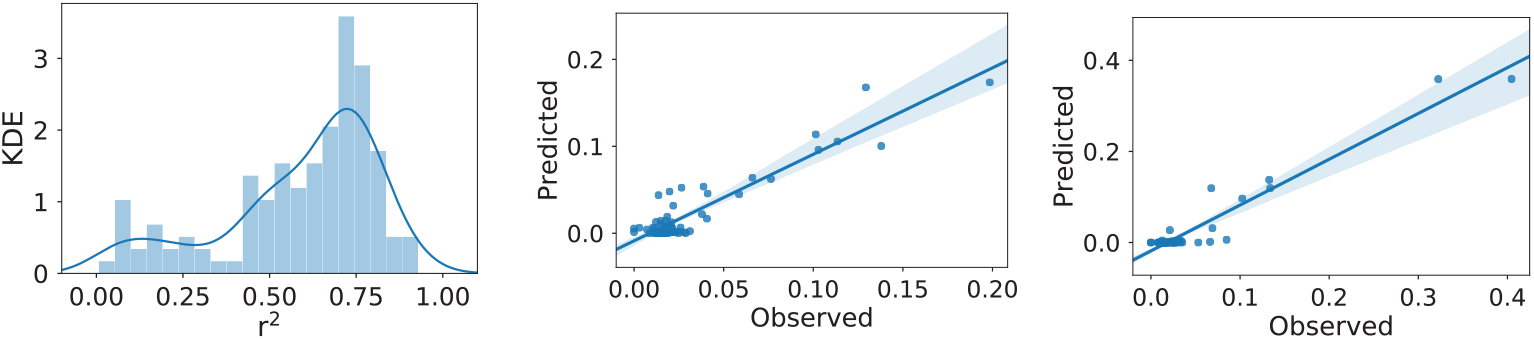
Excellent model performance in CN-individuals. (Left) The distribution of r^2^ values representing the range in individual-level model fit across all CN-subjects. Two exemplary subjects are plotted: (middle) a subject with very low tau burden; (right) a subject with low tau burden. Even at very low (subthreshold) levels, the distribution of tau is predicted by functional connectivity patterns.

### 3.4. Regional β-Amyloid is associated with region model performance

For each model, regions-of-interest were classified as either overestimated or underestimated by the model based on the sign of the residual (Fig 6A,B). Underestimated regions are those demonstrating greater tau burden than would be expected given connectivity to the model epicenter (i.e. observed > predicted), while overestimated regions demonstrate less tau than would be expected given their connectivity profile (i.e. predicted > observed). Compared to overestimated regions, underestimated regions had greater global *β*-amyloid burden (t = 3.72, p = 0.0004; Fig 6C,D), suggesting the regional presence of amyloid may accelerate the spread or expression of tau tangles. This effect was only present in amyloid+ individuals (Fig 6E).

**Figure 6:**
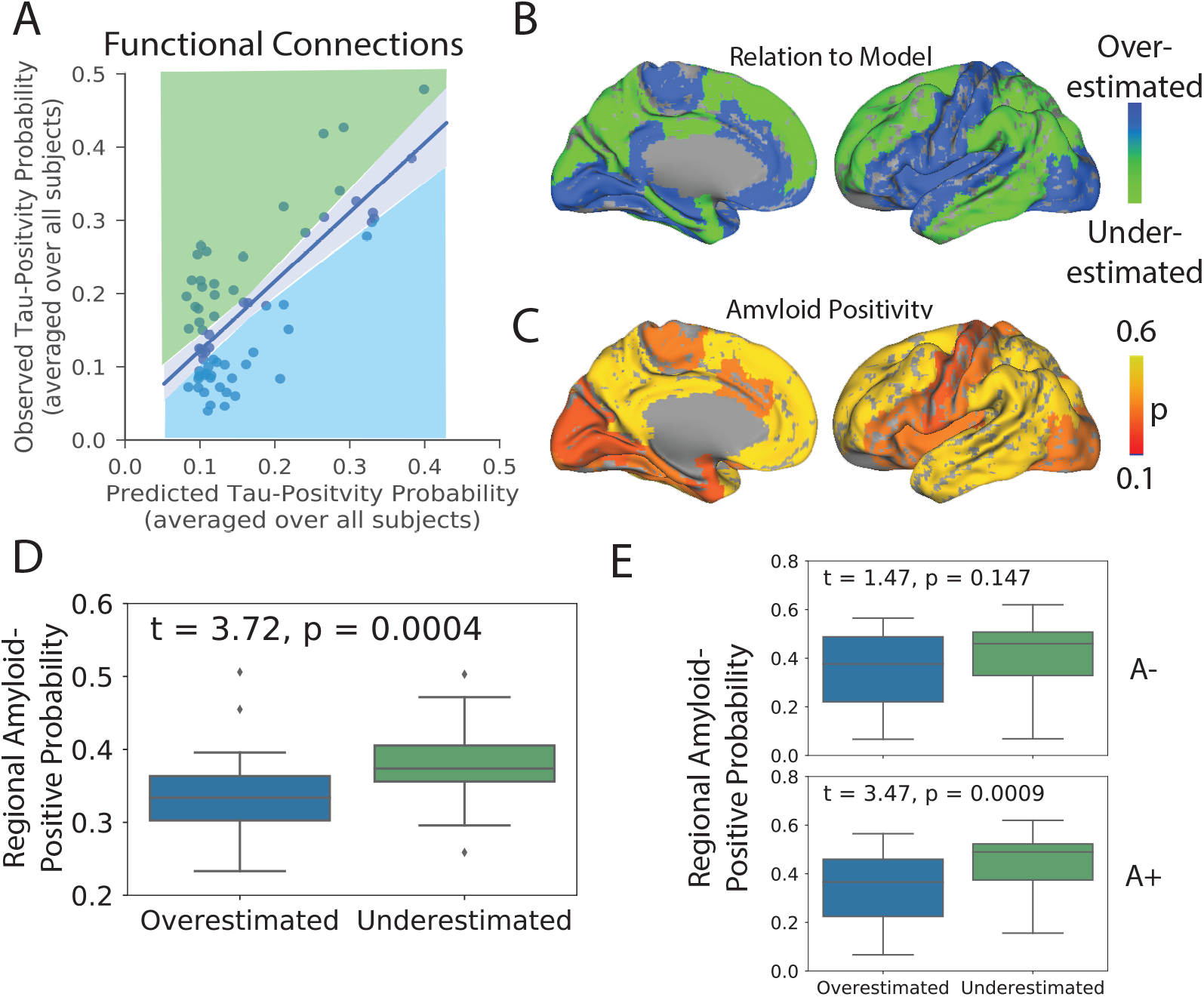
Amyloid explains regional model underestimation. A) Regions were classified as overestimated or underestimated based on the sign of the residual in a comparison of predicted vs. observed values. B) A surface render showing the spatial distribution of over- and underestimated regions. C) A surface render showing the spatial distribution of regional amyloid-positive probabilities averaged over all subjects. D) Underestimated regions tended to have significantly greater amyloid burden, suggesting these regions had more tau than would be predicted given their connectivity to the model epicenter. E) The same relationship stratified by amyloid status. A+ = Amyloid Positive; A-= Amyloid Negative

## 4. Discussion

Observations in post-mortem human brains [25, 24] and experiments in animal models [20, 21, 22, 23, 12] have together provided evidence that tau can be transmitted from cell to cell through neuronal projections. However, post-mortem studies cannot provide direct evidence of cell-to-cell spread, and while animal models have proven tau can spread through neuronal connections under certain unnatural conditions, they cannot prove that this phenomenon occurs naturally in humans. Studies searching for evidence of cell-to-cell transmission of tau in living humans have been limited by small datasets, simplistic models and issues relating to the quantitative measurement of tau. Here, we used a mixture-modeling approach on a large sample of humans on the Alzheimer’s disease spectrum to enhance the quantification of tau signal, and we applied to this data a diffusion model based on theoretical principles of an agent propagating through a network. These simulations explained a majority of the variance in the global spatial distribution of tau-PET signal in the brain, and performed nearly equally well in predicting the distribution of tau-PET signal in individual subjects. A similar model testing the hypothesis that tau spreads across neighboring brain regions was less successful at explaining the overall pattern. The models performed best in amyloid-negative individuals, and also systematically underestimated the magnitude of tau in regions classically shown to harbor *β*-amyloid. Together, these results suggest that tau spreads slowly through the limbic network in normal aging, and that the presence of *β*-amyloid leads to acceleration of tau tangle expression into isocortical regions.

Brain networks may be key to the evolution of neurodegenerative disease [45]. The atrophy patterns of many neurodegenerative dementias have been shown to resemble resting-state functional brain networks [46, 47], and network “hubs” are especially vulnerable to neurodegeneration across brain disorders [48]. Studies modeling the diffusion of gray matter degeneration across brain networks have recreated such patterns with impressive accuracy [47, 49, 50]. However, in many neurodegenerative disorders, brain atrophy is preceded and perhaps caused by the aggregation of pathological agents. In Alzheimer’s disease, the presence of tau is closely linked to [7, 8], and likely precedes [8, 11], gray matter atrophy. However, because gray matter degeneration observed in Alzheimer’s dementia may be caused by many sources other than Alzheimer’s pathology, gray matter degeneration itself cannot be used as proxy for tau (e.g. [51]). PET studies therefore provide a unique advantage by measuring pathological proteins more directly, and applying network diffusion models to PET data has, for example, led to the successful description of the spatial progression of *β*-amyloid in Alzheimer’s disease [43]. Our model uses a similar framework to simulate the spread of tau through the brain and reaches a similar level of success, both within-subject as well as globally across all subjects. The application of network models to other forms of dementia will be needed to conclude whether the spread of pathological proteins through connected neurons is a common thread linking many diseases.

While our model recapitulated the early stages of tau spreading accurately, later stages were modeled less accurately, with a systematic under-estimation of tau in regions prone to early and high-volume *β*-amyloid aggregation. While tau, not *β*-amyloid, is closely associated with atrophy in Alzheimer’s disease, the commonly-observed concurrence of extra-limbic tau and cortical amyloid burden has led to speculation that *β*-amyloid may accelerate or otherwise facilitate the spread of tau outside the medial temporal lobe. Recent studies in mice have shown that *β*-amyloid creates an environment facilitating the rapid fibrilization of tau [12, 13]. Our data support this notion, as brain regions harboring more *β*-amyloid, such as the precuneus and temporoparietal regions, had a higher incidence of abnormal tau than would be predicted simply by their regional connectivity to the medial temporal lobe. Further supporting this conclusion was the observation that this effect was only seen in amyloid-positive individuals. A conclusive model of tau spreading may not be complete without incorporating dynamic interaction with *β*-amyloid.

Tau tangles are a pathological hallmark of AD, but they are neither specific to AD, nor to neurodegenerative disease in general. The process of aging appears to lead inevitably to the accumulation of tau tangles in the medial temporal lobe and occasionally beyond, a phenomenon known as primary age-related tauopathy (PART) [9], and *in vivo* evidence for the longitudinal accumulation of tangles in healthy elderly has been observed [11]. While PART may result in subtle insults to cognition and brain health [52], there is still debate as to whether PART and AD are distinct processes [53]. We show that even in individuals without significant amyloid burden and low (subthreshold) tau-PET signal, the spatial pattern of tau can be predicted by functional connectivity to medial temporal lobe structures. These findings suggest that, even in PART, tau likely spreads from cell to cell through communicating neurons. The results also suggest closer scrutiny of subthresh-old tau-PET signal in cognitively unimpaired, amyloid-negative individuals. Elevated SUVR values occurring in a consistent pattern in specific limbic regions may be indicative of very low tau pathology, rather than non-specific or off-target ligand binding.

While our findings lend strong support to the hypothesis of tau spreading through communicating neurons, connectivity patterns and regional amyloid burden together could not fully explain the observed pattern of tau-PET across the brain. While a portion of this discrepancy may be explained by measurement error, there are likely other factors at play. Recent work has outlined a consistent genomic profile across regions that express tau [54], implicating regional variation in intrinsic molecular environment may mediate the presence and rate of tau tangle formation. This may explain why, for example, many subcortical regions do not show substantial tau burden despite connections to regions expressing neurofibrillary tau tangles. In addition, it is also possible that only certain neuron types can facilitate the transmission of tau, which may be challenging to model using macroscopic measures of functional connectivity. Finally, some studies have suggested the directional flow of neuronal activity may influence the spread of brain pathology [55]. Future studies incorporating this information, along with dynamics related to regional amyloid burden and regional vulnerability, may achieve a more complete model of tau spreading. However, at present, we show that the spread of tau is predicted by connectivity patterns to a degree that greatly exceeds both chance and other hypotheses of tau spread, and does so in a parsimonious fashion, greatly supporting the notion that connectivity is in some way involved in the spread of tau through the human brain.

Tau-PET signal has been notoriously hard to analyze due to extensive off-target binding reducing signal-to-noise ratio (for review, see [27]). We partially circumvented this well-known issue by applying Gaussian mixture-models separately to each region-of-interest. This approach effectively established a region-specific baseline representing the normal distribution of off-target signal, allowing the identification of outliers expressing SUVR values exceeding the normal expected range. A similar approach has been applied to the spatial staging of brain amyloid, leading to results that were highly consistent across samples [38]. However, this approach used a single threshold for all regions, whereas our approach was executed separately across each region, thereby accounting for regional ligand dynamics. The conversion of tau-PET SUVR values to tau-postive probabilites resulted in a clean distribution of values across the brain that greatly resembled the progressive pattern described in the pathology literature, and validated the expectation of no substantial burden in the striatum. By both treating each ROI separately but also expressing values along a standardized 0-1 probability scale, we were able to achieve greater regional sensitivity for the detection of both low-level tau, as well as high confidence tangle aggregation. Importantly, this approach did not require any arbitrary threshold (e.g. [56]) and resulted in discreet probability values, and therefore may benefit future studies or clinical evaluations seeking to classify regions as “tau-positive” with a given level of confidence.

Our study comes with a number of limitations. The premise of testing the hypothesis of tau spread through communicating neurons requires that both neuronal connections and tau burden are accurately measured. We attempt to partially surmount these issues by introducing a data-driven approach for overcoming off-target and non-specific binding in AV1451-PET data, and by validating our findings over different connectomes across different samples and modalities. Our mixture-modeling strategy is sensitive to sample size and composition. While it is unlikely that this phenomenon strongly affected the present findings, it is an important point worth consideration for future studies utilizing this approach to transform tau-PET data. Another limitation is raised by our choice to remove regions that do not demonstrate measurable tau burden, namely subcortical regions, from the model altogether. Certain subnuclei of subcortical structures such as the thalamus do accumulate tau pathology in Alzheimer’s disease [57], though we were unable to detect such pathology, perhaps due to the resolution of our measurements. While it is possible that subcortical structures participate in neuronal transmission of pathology without expressing the pathology itself, the current implementation of our model does not support this type of dynamic. However, while incidental measurement of indirect functional connectivity is a common critique of functional MRI, here it may pose an advantage, as functional connectivity mediated by subcortical connections may still be present in functional connectomes used for this study.

## 5. Conclusion

Altogether, our data strongly supports the notion that tau pathology itself, or information leading to the the expression of pathology, is transmitted from cell to cell in humans, principally through neuronal connections, and not extracellular space. Our findings further suggest that this phenomenon proceeds slowly but perhaps ubiquitously in normal aging, and that the process is accelerated dramatically in specific brain regions demonstrating *β*-amyloid burden. While our *in vivo* results cannot prove that tau spreads through neuronal connections, we show that more highly connected regions have a higher tendency to be affected closer in time by tau along a specific network path cascading from the medial temporal lobe. Future models may be able to improve results by incorporating region-specific vulnerability factors, directional flow and amyloid dynamics, though contributing such information in a parsimonious way presents a difficult challenge.

## Conflicts of Interest

OH has acquired research support (for the institution) from Roche, GE Healthcare, Biogen, AVID Radiopharmaceuticals, Fujirebio, and Euroimmun. In the past 2 years, he has received consultancy/speaker fees (paid to the institution) from Biogen, Roche, and Fujirebio.

## Acknowledgments

We would like to thank Bratislav Misic, Pierre Bellec, and Mallar Chakravarty for comments and suggestions during the formulation of this work, and and Liza Levitis for proofreading the manuscript. JWV is supported by government of Canada through the tri-council Vanier Canada Graduate Doctoral Fellowship. We would also like to acknowledge support from the Ludmer Centre for Neuroinformatics and Mental Health and the Healthy Brains for Healthy Lives initiative. Work at the authors research center was supported by the European Research Council, the Swedish Research Council, the Knut and Alice Wallenberg foundation, the Marianne and Marcus Wallenberg foundation, the Strategic Research Area MultiPark (Multidisciplinary Research in Parkinsons disease) at Lund University, the Swedish Alzheimer Foundation, the Swedish Brain Foundation, The Parkinson foundation of Sweden, The Parkinson Research Foundation, the Skne University Hospital Foundation, and the Swedish federal government under the ALF agreement. Doses of 18F-flutemetamol injection were sponsored by GE Healthcare. The precursor of 18F-flortaucipir was provided by AVID radiopharmaceuticals. Data collection and sharing for this project was funded by the Alzheimer’s Disease Neuroimaging Initiative (ADNI) (National Institutes of Health Grant U01 AG024904) and DOD ADNI (Department of Defense award number W81XWH-12-2-0012). ADNI is funded by the National Institute on Aging, the National Institute of Biomedical Imaging and Bioengineering, and through generous contributions from the following: AbbVie, Alzheimer’s Association; Alzheimer’s Drug Discovery Foundation; Araclon Biotech; BioClinica, Inc.; Biogen; Bristol-Myers Squibb Company; CereSpir, Inc.; Cogstate; Eisai Inc.; Elan Pharmaceuticals, Inc.; Eli Lilly and Company; EuroImmun; F. Hoffmann-La Roche Ltd and its affiliated company Genentech, Inc.; Fujirebio; GE Healthcare; IXICO Ltd.;Janssen Alzheimer Immunotherapy Research & Development, LLC.; Johnson & Johnson Pharmaceutical Research & Development LLC.; Lumosity; Lundbeck; Merck & Co., Inc.;Meso Scale Diagnostics, LLC.; NeuroRx Research; Neurotrack Technologies; Novartis Pharmaceuticals Corporation; Pfizer Inc.; Piramal Imaging; Servier; Takeda Pharmaceutical Company; and Transition Therapeutics. The Canadian Institutes of Health Research is providing funds to support ADNI clinical sites in Canada. Private sector contributions are facilitated by the Foundation for the National Institutes of Health (www.fnih.org). The grantee organization is the Northern California Institute for Research and Education, and the study is coordinated by the Alzheimer’s Therapeutic Research Institute at the University of Southern California. ADNI data are disseminated by the Laboratory for Neuro Imaging at the University of Southern California.

## Notes

#### Summary of Updates

The major change in this version was that we added more amyloid-individual (mostly MCI-) and tried to emphasize more that finding that spread is predicted even in amyloid negative individuals. All figures were revised except figure 1 to accomodate the addition of ~20 subjects.

